# Mapping inter-individual functional connectivity variability in TMS targets for major depressive disorder

**DOI:** 10.1101/2021.07.15.452518

**Authors:** Shreyas Harita, Davide Momi, Frank Mazza, John D. Griffiths

## Abstract

Transcranial magnetic stimulation (TMS) is an emerging alternative to existing treatments for major depressive disorder (MDD). The effects of TMS on both brain physiology and therapeutic outcomes are known to be highly variable from subject to subject, however. Proposed reasons for this variability include individual differences in neurophysiology, in cortical geometry, and in brain connectivity. Standard approaches to TMS target site definition tend to focus on coordinates or landmarks within the individual brain regions implicated in MDD, such as the dorsolateral prefrontal cortex (dlPFC) and orbitofrontal cortex (OFC). Additionally considering the network connectivity of these sites has the potential to improve subject-specificity of TMS targeting and, in turn, improve treatment outcomes. We looked at the functional connectivity (FC) of dlPFC and OFC TMS targets, based on induced electrical field (E-field) maps, estimated using the SimNIBS library. We generated individualized E-field maps on the cortical surface for 121 subjects from the Human Connectome Project database using tetrahedral head models generated from T1-weighted MR images. We analyzed inter-subject variability in the shape and location of these TMS target E-field patterns, their FC, and the major functional networks to which they belong. Our results revealed the key differences in TMS target FC between the dlPFC and OFC, and also how this connectivity varies across subjects. Three major functional networks were targeted across the dlPFC and OFC: the ventral attention, fronto-parietal and default-mode networks in the dlPFC, and the fronto-parietal and default mode networks in the OFC. Inter-subject variability in cortical geometry and in FC was high. Our results characterize the FC patterns of canonical therapeutic TMS targets, and the key dimensions of their variability across subjects. The high inter-individual variability in cortical geometry and FC, leading to high variability in distributions of targeted brain networks, may account for the high levels of variability in physiological and therapeutic TMS outcomes. These insights should, we hope, prove useful as part of the broader effort by the psychiatry, neurology, and neuroimaging communities to help improve and refine TMS therapy, through a better understanding of the technology and its neurophysiological effects.

**Highlights:**

- E-field modelling and functional connectivity used to study TMS targets (dlPFC,OFC)
- Considerable variability in TMS target E-field patterns seen across subjects
- Large inter-subject differences in target connectivity observed and characterized
- Major functional networks targeted by dlPFC, OFC TMS were the VAN, FPN and DMN
- Insights can contribute to improved and more personalized TMS therapies in the future

## Introduction

### TMS stimulation therapy targets and the neurobiology of major depressive disorder

A considerable number of patients with major depressive disorder (MDD) do not respond to first-line therapies such as drugs or psychotherapy (MacQueen et al., 2017). People who fail two or more pharmacological interventions of a sufficient dose and time are characterized as having treatment-resistant depression (TRD; (Souery et al., 1999). Transcranial magnetic stimulation (TMS) is an efficacious and cost-effective treatment for people with TRD and is an emerging alternative to existing treatments for MDD, as well as a variety of other neurological and psychiatric disorders. However, the clinical utility of TMS remains limited by the large heterogeneity in its clinical outcomes (Nestor & Blumberger, 2020). One factor believed to contribute to this variable clinical response among patients is individual differences in structural and functional brain connectivity (Downar & Daskalakis, 2013). In order to find a target site for TMS treatment, seed-based approaches have focused on individual brain regions implicated in MDD, such as the dorsolateral prefrontal cortex (dlPFC), dorsomedial prefrontal cortex (dmPFC), and orbitofrontal cortex (OFC). On the other hand, network-based approaches have shown how TMS efficacy can be improved by considering not only the location of the primary stimulation site (dlPFC, OFC, etc.), but also its connectivity - i.e. the wider set of distal brain regions that are mono- or poly-synaptically activated by TMS stimulation (Drysdale et al., 2017; Fox et al., 2013).

Previous studies have shown that variability in clinical efficacy of dlPFC-targeted repetitive TMS (rTMS) treatment for MDD is related to differences in the functional connectivity (FC) of the specific dlPFC locations stimulated (see Fox et al., 2012, Cash et al., 2020). These observations suggest that a detailed examination of individual differences in FC patterns for frontal lobe rTMS targets should prove useful in further refining TMS targeting methodologies. The aim of the present study was to undertake such an examination. Specifically, we characterized the FC patterns of the dlPFC and OFC, based on E-field maps generated by biophysical simulations of TMS stimulation effects on the cortex (see *Methods*). Our hypothesis was that individual differences in spontaneous functional brain dynamics would contribute more to downstream network engagement than individual differences in cortical geometry (i.e., E-field variability), which may, in turn, explain some of the observed heterogeneity of rTMS treatment outcomes.

By far the most commonly targeted site in rTMS treatment of MDD is the dlPFC. That this region is known to play a critical role in executive functions such as attention, planning, and organization, and up-regulation of the circuits underlying these neurocognitive functions is one potential explanation for its positive therapeutic effects. rTMS stimulation of the left dlPFC has also been observed to regulate FC to and between the reward- and emotion-related regions of the mesocorticolimbic dopamine pathway, which have also been consistently implicated in MDD (Tik et al., 2017). However, stimulation of the left dlPFC does not work for all patients, with only around 46% of MDD patients achieving response, and only 31% achieving remission after a standard rTMS treatment course (Fitzgerald et al., 2016). This may be due to inter-subject differences in neurochemistry, connectivity, in their specific MDD neuropathology, or a variety of other potential factors. To overcome this issue of limited success with dlPFC targeting, several groups have begun to explore alternative rTMS targets to treat MDD, such as the dmPFC and the OFC (Downar & Daskalakis, 2013).

Recent research has shown that the OFC is hyperactive in MDD (Feffer et al., 2018). The OFC consists of a medial (mOFC) and a lateral (lOFC) subdivision, each with unique anatomical and FC profiles. The OFC has extensive cortico-cortical and cortico-striatal connections to regions implicated in MDD such as the cingulate cortex, caudate, striatum, hypothalamus, amygdala, hippocampus, insula, and thalamus (Zald et al., 2014). Recently, studies have begun to explore the OFC as a TMS target, with the rationale being to stimulate these MDD-implicated cortico-cortical and cortico-striatal loops. The OFC and its downstream connections are principal contributors to reward and reversal learning (Kringelbach, 2005), predictive and fictive error assessment (Boorman et al., 2013), emotional regulation, and generation of affective states (Rolls, 2019). Due to the wide psychiatric implications of pathological OFC activity, this region has been growing in popularity as an alternative rTMS treatment target for mental illness. For example, low frequency (1Hz) rTMS of the left OFC has been shown to ameliorate symptoms in patients with obsessive-compulsive disorder (OCD; (Kumar et al., 2018). Relatedly, 1Hz rTMS to the right OFC in a recent study saw nearly a quarter of patients suffering from MDD achieve remission (Feffer et al., 2018). Importantly, these patients previously showed minimal response to dmPFC-rTMS. These results point to heterogeneous mechanisms of action, and therefore therapeutic effect, for dlPFC-, dmPFC-, and OFC-rTMS, possibly due to their unique downstream connections.

### E-field modelling and cortical geometry

TMS uses high-intensity magnetic field pulses to influence neuronal activity. The TMS coil produces a time-varying magnetic field, which in turn induces a focal electric field (E-field) within brain (principally cortical) tissue. E-field modelling is a relatively new approach that uses computationally estimated E-field maps, which can serve as a proxy to identify the region of the brain that is stimulated for a given coil type, location, orientation, and (MRI scan-derived, subject-specific) head and brain characteristics (Opitz et al., 2011; Thielscher et al., 2015; Weise et al., 2020). These methodologies are increasingly used in clinical and basic TMS research, as a means to better understand and minimize the sources of variability in TMS outcomes due to the varying placement of TMS coils on the subject’s scalp, and to variability in each individual’s skull anatomy and cortical geometry.

Approaches to TMS coil placement include the ‘5cm-rule’, 10-20 EEG electrode locations, MRI-guided anatomical targeting, and the more recent fMRI FC-guided targeting. However, there is a considerable amount of variability in the ‘ideal’ TMS coil placement to optimally stimulate a specific target. For example, the F3 10-20 electrode position may result in different parts of the dlPFC being stimulated in different individuals. Moreover, previous research has shown that differences in the complex neuroanatomy of each individual human skull and brain (i.e., brain size, gyri, and sulci differences) result in E-fields of varying shapes, sizes, and depths (Thielscher et al., 2011). The combined effect of TMS coil placements and subject-specific differences in skull anatomy and cortical geometry leads to an inter-subject variation in E-field patterns across patients/subjects and is a potential explanation for some of the high variability in rTMS therapy outcomes. Furthermore, the interaction between this cortical geometry-driven variability in TMS-induced E-fields and variability in intrinsic FC patterns, despite receiving some attention from researchers previously (Opitz et al., 2016), remains poorly understood.

### Present study

It is likely that the dlPFC and OFC have unique functional connections, through which they each exert their differential therapeutic effects following rTMS treatment (Vila-Rodriguez & Frangou, 2021). The patterns of these connections likely vary considerably across subjects, and very little is currently known about the relative contributions of these variability sources to TMS treatment outcomes. In this framework, even though a previous study has combined anatomically realistic finite element models of the human head with resting functional MRI to predict functional networks targeted via TMS at a given coil location and orientation (Opitz et al., 2016), none so far have investigated the impact of the individual subject and group-average FC patterns for a consistent E-field-defined seed region using different TMS locations. In the present study, we, therefore, sought to address part of this knowledge deficit, by systematically examining E-field and FC patterns for dlPFC and OFC TMS target sites, using structural and functional neuroimaging scans in a group of healthy control subjects. We computed simulated TMS E-fields centered at the dlPFC and OFC and studied their FC patterns using both individual and group averaged resting-state fMRI data. We found specific networks being targeted as a result of dlPFC or OFC TMS. While the same major networks were targeted consistently across our subject cohort, substantial inter-individual differences in each subject’s specific relation to these networks were also observed. Comparing individual subject and group-average FC patterns for a consistent E-field-defined seed region allowed us to assess the contribution of inter-individual variability in FC patterns, independently of variability in cortical geometry. Conversely, comparing FC patterns for a single group-level E-field seed to those for subject-specific E-field seeds allowed us to quantify variability in TMS target connectivity due purely to skull and cortical geometry variation. Our results highlight the inter-individual differences in dlPFC and OFC TMS FC, potentially paving the way for personalized rTMS therapy in the future.

## Methods

Analysis of T1-weighted anatomical MRI (for E-fields) and fMRI (for FC) data was conducted for 121 randomly selected subjects from the human connectome project (HCP) database, looking specifically at FC patterns related to dlPFC and OFC TMS stimulation. We computed TMS E-fields using the SimNIBS software package and focused on the cortical surface component of the stimulated tissue. The group average and individual subject HCP CIFTI dense connectomes were used to determine the FC of TMS targets to the rest of the brain, and we used the standard Yeo/Schaefer parcellations to summarize the downstream connections of the dlPFC and OFC. Finally, we defined a framework for assessing contributions to overall variability from cortical geometry, FC, and from a combination of these two sources. Each of these steps is detailed below.

### Determining the TMS E-field

The TMS-induced electric field was modelled using tools from the SimNIBS software library (Thielscher et al., 2015). We used the ‘mri2mesh’ head modelling pipeline to create a tetrahedral surface mesh head model (.msh file) from T1-weighted MR images and Freesurfer tissue segmentations for all subjects. This mesh consisted of five tissue types: white matter (WM), grey matter (GM), cerebrospinal fluid (CSF), skull, and scalp. The assigned conductivity values were fixed, as per the SimNIBS defaults: 0.126 S/m (WM), 0.275 S/m (GM), 1.654 S/m (CSF), 0.01 S/m (skull), and 0.465 S/m (scalp). In order to investigate possible effects due to head geometry, two head mesh types were used as part of the SimNIBS analysis pipeline. The first was each subject’s unique head meshes, as derived from that subject’s own neuroanatomical MRI scans. The second was the general template head mesh, ‘ernie.msh’, which is distributed as a part of SimNIBS. The EEG 10-20 system F3 electrode was selected to target the left dlPFC, for two reasons: First: EEG F3 is in our experience currently the most commonly used left dlPFC targeting method in clinical rTMS practices. Second: it has been reported that TMS targeting approaches based on the 10-20 EEG system account better for variability across different skull shapes and sizes than scalp-based measurements such as the ‘5cm rule’ (Cash et al., 2020). With regards to the OFC, previous work has shown that targeting the right OFC via the Fp2 electrode led to remission in MDD patients unresponsive to dlPFC- and dmPFC-rTMS (Fettes, 2020). Given the high level of anatomical symmetry between hemispheric homologues, here we used the Fp1 electrode (left-side homologue of Fp2, thus targeting left OFC), so as to keep both TMS targets in the left hemisphere. This approach enabled us to make more direct comparisons between dlPFC and OFC, minimizing extraneous methodological differences. We strongly expect our left OFC results to generalize well to right OFC targets, although we leave the full demonstration of this for future work. The left dlPFC has been used as a target for rTMS therapy almost since the technique’s inception (George et al., 1995), and while there are heterogeneous outcomes associated with left dlPFC rTMS, it is still one of the most widely used rTMS targets for MDD (Cash et al., 2020; Pascual-Leone et al., 1996). The use of the OFC as a TMS target to treat psychiatric disorders, while still a novel and largely underexplored idea, has recently gained traction - with OFC rTMS showing promise in treating MDD (right OFC; Feffer et al., 2018) and OCD (left OFC; Kumar et al., 2018).

At both coil centers (F3, Fp1), the coils were positioned using the standard orientation to ensure that the resulting E-field is directed perpendicularly into the cortex. Previous studies have shown that this standard orientation is able to achieve the highest perpendicular E-field values (Janssen et al., 2015). This is done by pointing the coil handle away from the midline of the cortex. The y-direction position values for the dlPFC (F3) and OFC (Fp1) are therefore F5 and AF7, respectively. Of the various coil models available in SimNIBS, we used the Magstim 70mm Figure-8 coil, which is the most common coil type in both clinical and research settings. To keep our focus primarily around the target regions (dlPFC, OFC), we used a threshold of 0.9 Volt/meter (V/m) to limit the size of the E-field obtained (Romero et al., 2019). We report the E-field sizes in units millimeter squared (mm^2^), calculated directly from the faces (triangles) of the CIFTI surface meshes. For reference, these meshes consist of 32,492 vertices and 64,980 faces per hemisphere, with an average face area of 0.05 mm^2^.

### Functional Connectivity

Resting-state fMRI data of the 121 HCP subjects was used to study the FC of TMS target regions. For full details on the HCP acquisition protocols and related information, see (Glasser et al., 2013; Uğurbil et al., 2013; Van Essen et al., 2013;. Van Essen et al., 2012).

For subject-specific FC analyses, the FC for the CIFTI format time series for each HCP subject’s four resting-state fMRI scans were averaged and converted into ‘dense connectome’ (Pearson correlation) FC matrices, each containing 91,282 rows and columns (corresponding to ∼64,000 cortical surface vertices, and ∼27,000 sub-cortical voxels). For group-level FC analyses, the HCP_S1200_GroupAvg_v1 (1003 subjects) dense connectome was used instead of individual-subject data. The FC of a given E-field was determined by taking the average FC, over all the vertices within that E-field, to every other node in the dense connectome FC matrix. Note that in this study we only studied connectivity within the stimulated (i.e. the left) hemisphere. In order to summarize which downstream regions were functionally connected to the stimulated areas, we grouped the connectivity profiles of dlPFC and OFC stimulation target sites according to the canonical functional network parcellation of Yeo et al. (2011) and Schaefer et al. (2018). These canonical networks consisted of the visual (Vis), somatomotor (SomMot), dorsal attention (DAN), ventral attention (VAN), limbic, frontoparietal (FPN), temporoparietal (TempPar), and default-mode (DMN) networks (Yeo et al., 2011, Schaefer et al., 2018). These canonical Yeo/Schaefer network summaries give a useful low-dimensional complement to the high-dimensional (E-field seed column-averaged) FC dense connectome columns, helping us to gain a better understanding of which functional networks might be stimulated by TMS, and how the pattern of stimulated areas varied between target sites (dlPFC, OFC) and across subjects. Here, the individual units of a Yeo/Schaefer parcellation-based functional connectome are the brain regions identified by (Schaefer et al., 2018), and serve as building blocks for functional brain anatomy. In the context of network analysis, each parcel represents a single node within a whole-brain network. In the following, we, therefore, refer to these individual Yeo/Schaefer parcels as network ‘nodes’. In all subjects, we analyzed E-field variability, FC patterns, and the major nodes of the most common functional networks and the FC maps they created.

To explore the impact of individual brain features on the variability of TMS target connectivity, we delineated a two-level FC analysis framework. At the first level (1a, 1b), the two most likely main sources of TMS FC variability are analyzed separately, and at the second level (2) they are analyzed in combination:

*1a.* *Influence of individual head, skull, and cortical geometry on TMS target connectivity patterns*. To examine this we computed the variability shown in the E-fields by holding the FC constant. To do this, we used each subject’s unique head mesh to determine their individualized E-field map. As described above, the connectivity of each subject’s specific E-field to the canonical HCP_S1200_GroupAvg_v1 resting-state FC matrix was studied.
*1b.* *Inter-subject differences in TMS target connectivity due purely to each subject’s unique functional connectivity profile*. This line of analysis involved using the same E-field across all subjects, but combining it with individualized FC. The SimNIBS general-template head mesh (‘ernie.msh’) was used to generate a fixed E-field pattern for all subjects, for each of the two TMS targets. The connectivity patterns of this fixed E-field to the rest of the brain were calculated using each subject’s individual FC matrix, derived from their four resting-state fMRI scans. This approach allowed us to measure the effect of individual spatial FC fingerprints on TMS target connectivity.
*2.* *Combined influence of individual cortical geometry and individual functional connectivity structure on TMS target connectivity patterns*. To represent the ‘real-world’ scenario, where individual characteristics of both cortical geometry and FC jointly contribute to TMS target connectivity patterns, we combined the approaches in 1a and 1b above and studied patterns using both each subject’s unique head mesh and their specific FC matrices.

### Statistical Analysis

Connectivity scores were compared separately for dlPFC and OFC using repeated-measures one-way ANOVA, with the (within-subjects) factor “NETWORK” (8 levels for the 8 functional networks: Vis, SomMot, DAN, VAN, Limbic, FPN, TempPar, DMN). Subsequent pairwise post hoc comparisons were performed to determine significant differences between NETWORK levels. The critical p-value was then adjusted using Tukey correction to account for multiple comparisons (**.05; Tukey corrected; *.05 uncorrected).

### Code and Data Availability

All analyses reported in this paper were conducted on CentOS Linux compute servers running Python 3.7.3, using the standard scientific computing stack and several open-source neuroimaging software tools - principally SimNIBS (E-field simulations; (Thielscher et al., 2015), Nibabel (neuroimaging data I/O; (Brett et al., 2020) and Nilearn (neuroimaging data visualizations; (Abraham et al., 2014). All code and analysis results are openly available at github.com/griffithslab/HaritaEtAl2021_tms-efield-fc.

## Results

### Influence of individual cortical geometry on TMS target connectivity patterns

#### E-field variability

There was considerable variability in the size of the E-fields across the subject group, as defined by the spatial extent of the thresholded E-field surface maps. At the dlPFC, E-field size ranged from 12 to 112.5 mm^2^ (mean = 54.3 ± 18 mm^2^). The OFC on the other hand was smaller in terms of overall E-field size, ranging from 3.2 to 60.7 mm^2^ (mean = 16.2 ± 8.5 mm^2^). The E-field sizes varied to a greater extent for dlPFC stimulation (scalp position F3) than for OFC stimulation (scalp position Fp1) (Fig. 2 - ***Panel A***). (See *Methods* for information on the threshold value chosen and on physical dimensions of surface units).

**Figure 1:**
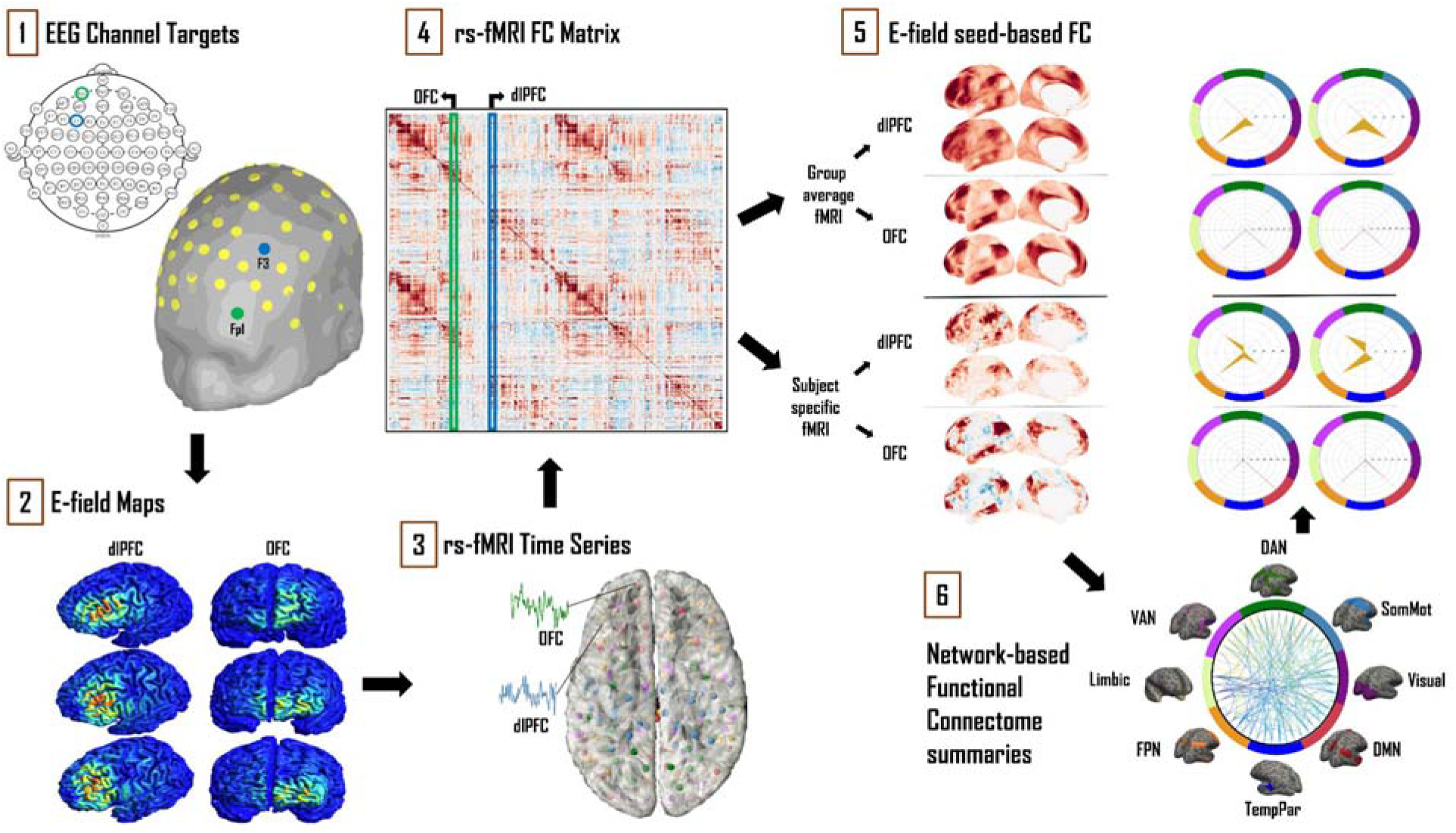
Schematic of the analytical approach. **1)** Left dlPFC and OFC 10-20 EEG electrode locations were identified as F3 and Fp1, respectively. **2)** SimNIBS simulations were run using these electrode locations as TMS coil placements, with the main output of interest being cortical surface E-field maps. **3)** Resting-state fMRI time series for 121 subjects from the HCP database were averaged and converted into ‘dense connectome’ FC matrices. **4)** The connectivity patterns of each subject’s E-field were determined using these dense connectomes. **5)** FC maps of the resulting E-fields were thus obtained with group-averaged and subject-specific resting-state fMRI data. **6)** These maps were further analyzed and summarized in terms of connectivity to the canonical multi-network parcellation templates of Yeo et al., (2011). The most prominent networks targeted by dlPFC and OFC TMS are reported. Spider plot visualizations in this example and later figures show the networks being targeted as a percentage (area of the orange polygon) of the suprathreshold E-field vertices.

**Figure 2:**
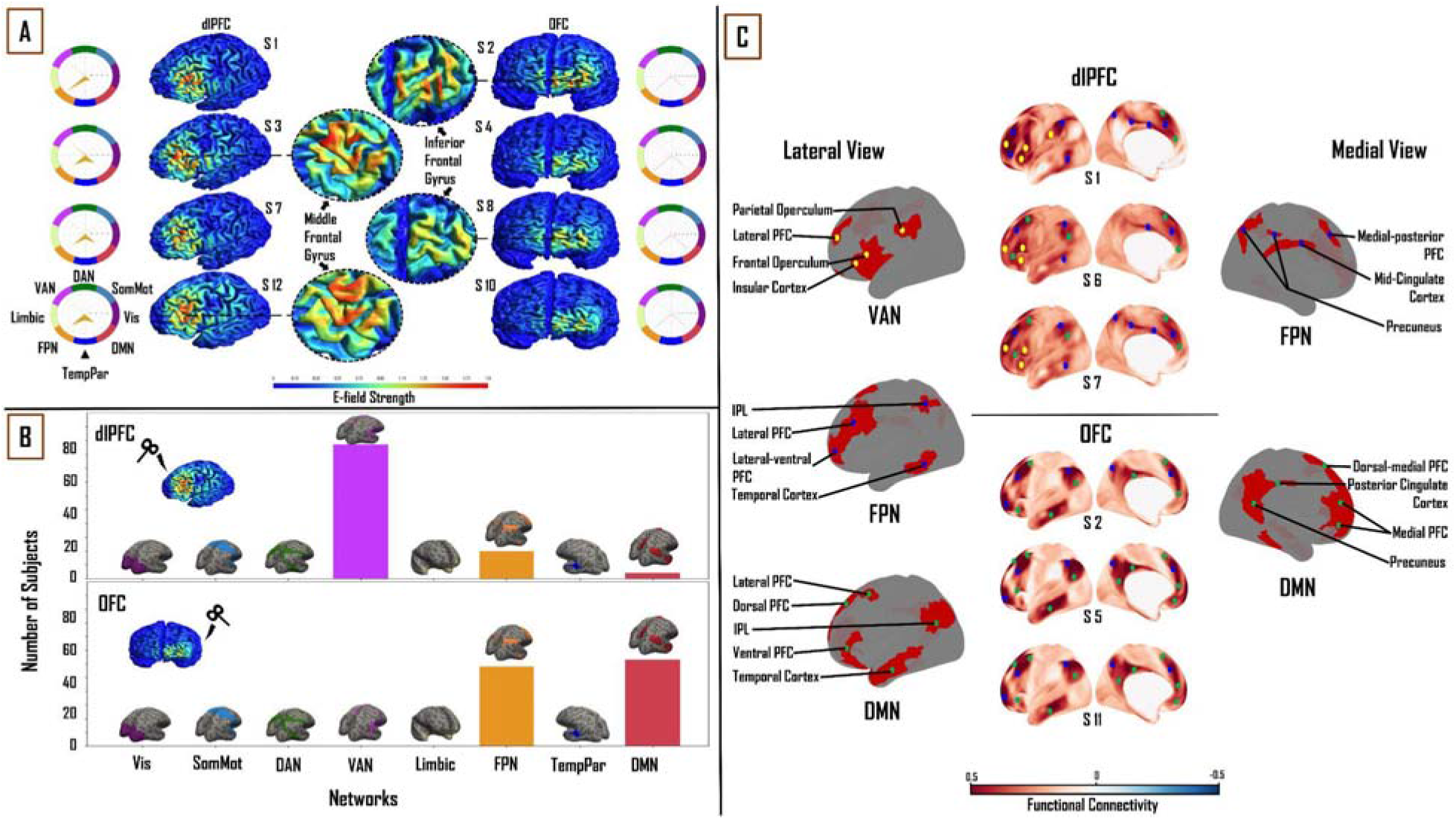
Influence of individual cortical geometry on TMS target connectivity. **<u>A)</u>** Individual subject E-fields for a subset of subjects highlighting anatomical differences between dlPFC and OFC E-fields. Spider plots on either side show functional network connectivity (expressed as a percentage of E-field vertices), based on the group-average FC matrix. E-field unit = V/m. **<u>B)</u>** Summary statistics of network engagement over all 121 subjects. Top: VAN, FPN, and DMN were the main networks engaged from the dlPFC. Bottom: FPN and DMN were the main networks engaged from the OFC. **<u>C)</u>** *Top:* Lateral and medial view of dlPFC FC maps in subjects 1, 6, and 7, highlighting the key regions that are functionally connected across VAN, FPN, and DMN. These regions lie mainly in frontal, parietal, and temporal cortices. *Bottom:* Lateral and medial view of OFC FC maps in subjects 2, 5, and 11, highlighting key regions that are functionally connected across FPN and DMN. These regions lie mainly in medial-frontal, cingulate, and posterior parietal cortices. VAN = ventral attention network, FPN = frontoparietal network, DMN = default-mode network.

#### Functional network connectivity based on subject-specific E-fields

We analyzed connectivity strength within each subject’s E-field maps by comparing maximum FC values (network engagement). A significant main effect of “NETWORK” was found at the dlPFC (F_(1,7)_= 1027.55, p < 0.0001, η2= 0.90) and OFC (F_(1,7)_= 1907.36, p < 0.0001, η2= 0.94). Across all subjects, three networks from the (Yeo et al., 2011) functional network parcellations had maximum FC to vertices in the dlPFC and OFC E-fields. In the dlPFC, the VAN (38.2 ± 8%), FPN (27.9 ± 7%), and DMN (20.3 ± 6.4%) accounted for an average of 86% of the E-field vertices. In the OFC, the FPN (46.5 ± 10.3%) and DMN (51.4 ± 10.3%) accounted for 98% of the E-Field vertices. The VAN, FPN and DMN were the most engaged networks from the dlPFC across the group (97, 20 and 4 out of 121 subjects, respectively). The DMN was the more engaged network from OFC than the FPN (63 and 58 out of 121 subjects, respectively). (Fig. 2 - ***Panel B***). Pairwise t-tests showed strong interactions between the TMS stimulation sites and the three major functional networks. We found the OFC target region showed greater connectivity to the DMN (T = 29.2; p <0.0001) and FPN (T = 19; p <0.0001) relative to the dlPFC target region, whereas the dlPFC target region showed greater connectivity to the VAN (T = 48.3; p <0.0001).

#### Relationship between TMS targets and downstream brain regions in subject-specific E-fields

After summarizing the overall structure of TMS target connectivity to the rest of the brain in terms of E-field vertex FC to the eight canonical Yeo networks, we examined more closely the spatial topographies of these FC patterns. Specifically, we studied the seed-based FC maps (where the seed is the entire thresholded E-field, and the maps are averaged over vertices within the seed) for each subject and target site, and identified through extensive manual inspection the dominant and consistent sub-patterns within those maps. Specific brain regions in the VAN, FPN, and DMN were highlighted with dlPFC-TMS stimulation. On the lateral cortical surface, key nodes within the VAN included the frontal and parietal opercula, lateral prefrontal cortex (PFC), and insular cortex. The main lateral FPN nodes were the posterior part of the middle and inferior temporal gyri, inferior parietal lobule (IPL), and the lateral-ventral and lateral PFC. Medial FPN nodes included the precuneus, mid-cingulate cortex, and medial-posterior PFC. Within the DMN, we observed the IPL, the lateral and ventral PFC on the lateral cortical surface; while the dorsal-medial and medial PFC constituted the medial DMN nodes. On the medial surface of the cortex, we observed FPN and DMN nodes, however, there were no specific VAN nodes (Fig. 2 - ***Panel C* [top]**). With regards to the OFC, specific regions in the FPN and DMN were highlighted. Within the FPN, laterally, we observed several of the same nodes noted above, including the lateral-ventral, lateral PFC, and IPL. Medial FPN nodes included the medial-posterior PFC and precuneus. In the DMN, the IPL and lateral PFC were seen once again as key nodes laterally. Medial DMN nodes included the dorsal-medial, medial PFC, and precuneus. DMN nodes specific to the OFC included the dorsal PFC and the anterior portion of the middle and inferior temporal gyri on the lateral cortical surface; and the posterior cingulate cortex (PCC) on the medial cortical surface (Fig. 2 - ***Panel C* [bottom]**). Critically, similar key nodes within different functional networks showed markedly different FC patterns between the two TMS target sites.

### Influence of connectivity structure on TMS target connectivity patterns

In the previous section, we held the FC matrix fixed, allowing us to characterize inter-subject differences in (putative) TMS target connectivity resulting purely from variation in the head, skull, and brain anatomy and geometry. We now examine the reverse scenario: inter-subject differences in TMS target connectivity due purely to the individualized FC, but using a single fixed E-field map for all subjects.

#### E-field Variability

The size of the constant E-field at the dlPFC was 310 mm^2^ and at the OFC was 6.3 mm^2^ (Fig. 3 - ***Panel A***).

**Figure 3:**
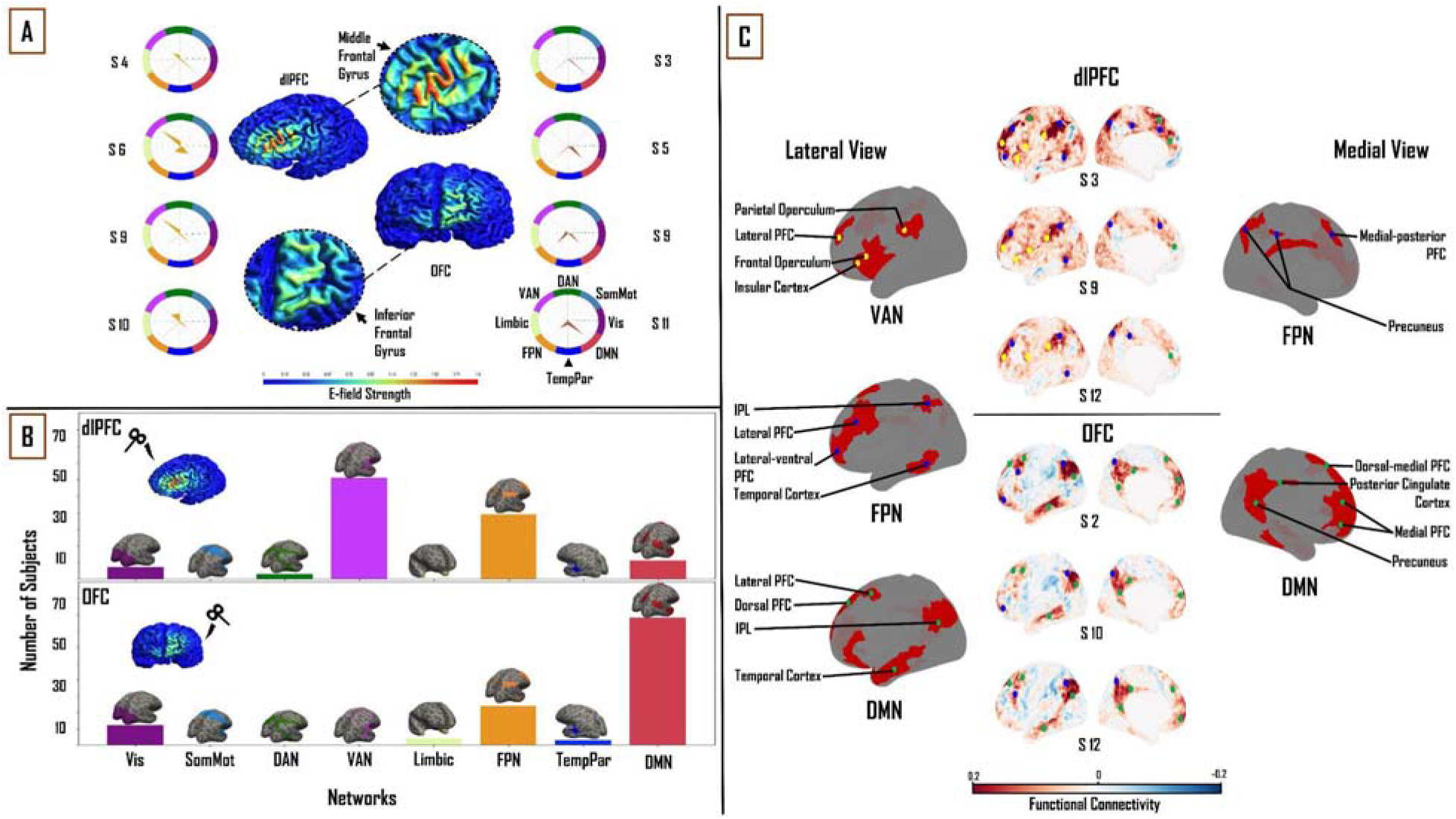
Influence of individual subject FC on TMS target connectivity. **<u>A)</u>** constant (Ernie-based) dlPFC and OFC E-fields, from F3 and Fp1 TMS targets, are 31 mm^2^ and 6.3 mm^2^ in size, respectively. Spider plots on either side show functional network connectivity (expressed as a percentage of E-field vertices), based on subject-specific FC matrices. E-field strength unit = V/m. **<u>B)</u>** Similar to Fig. 2, summary statistics of network engagement over all 121 subjects shows that the VAN, FPN, and DMN were the main networks engaged from the dlPFC [top]; while the FPN and DMN were the main networks engaged from the OFC [bottom]. **<u>C)</u>** *Top:* Lateral and medial view of dlPFC FC maps in subjects 3, 9, and 12, highlighting the key regions that are functionally connected across the VAN, FPN, and DMN. These regions lie mainly in frontal, parietal, and temporal cortices. *Bottom:* lateral and medial views of OFC FC maps in subjects 2, 10, and 12, highlighting key regions that are functionally connected across the FPN and DMN. These regions lie mainly in medial-frontal, cingulate, and posterior parietal cortices (bottom). VAN = ventral attention network, FPN = frontoparietal network, DMN = default-mode network.

#### Functional network connectivity based on constant E-fields

One-way ANOVA identified a significant main effect of “NETWORK” for both dlPFC (F_(1,7)_= 264.37, p < 0.0001, η2= 0.69) and OFC (F_(1,7)_= 208.05, p < 0.0001, η2 = 0.64). The same three functional networks from the subject-specific analysis were seen to have maximum FC to vertices in the constant E-field across subjects. In the dlPFC, dominant connectivity to the VAN (30.6 ± 12.3 %), FPN (29 ± 10.3%) and DMN (21.5 ± 7.9%) together accounted for 81% of E-field vertices. In the OFC, the FPN (23.4 ± 13.4%) and DMN (42 ± 16.3%) together accounted for 65% of the E-Field vertices. Once again, the VAN, FPN and DMN were the most engaged networks from the dlPFC across the group (61, 39 and 11 out of 121 subjects, respectively). The DMN was more engaged than the FPN in a majority of subjects, from the OFC (78 and 24 out of 121, respectively) (Fig. 3 - ***Panel B***). Once again, pairwise t-tests showed strong interactions between the TMS targets and the three major functional networks. We found the OFC target region showed greater connectivity to the DMN (T = 13.6; p <0.0001) relative to the dlPFC target region, whereas the dlPFC target region showed greater connectivity to the VAN (T = 25.2; p <0.0001) and FPN (T = 4.1; p<0.0001).

#### The relation between TMS targets and downstream brain regions in constant E-fields

Turning again to a detailed inspection of the seed-based FC maps, specific brain regions in the VAN, FPN, and DMN were highlighted with dlPFC-TMS, and within the FPN and DMN in OFC-TMS. Within the VAN, similar to our findings in the previous section, the lateral PFC, frontal and parietal opercula, and the IPL were the key nodes observed on the lateral cortical surface. The lateral FPN nodes included the posterior regions of the middle and inferior temporal gyri, the IPL, and the lateral-ventral and lateral PFC. We noted the precuneus and medial-posterior PFC as the main medial FPN nodes. Key DMN nodes included the lateral PFC and the dorsomedial and medial PFC. Again, as in the previous section, on the medial surface of the cortex, we observed FPN and DMN nodes, but no specific VAN nodes were seen (Fig. 3 - ***Panel C* [top]**). At the OFC, on the lateral left cortical surface, the FPN included some of the same nodes noted above such as the lateral-ventral and lateral PFC and the IPL. The main lateral DMN nodes included the IPL, the anterior portion of the middle and inferior temporal gyri, and the dorsolateral PFC. On the medial left cortical surface, the DMN nodes included the dorsal-medial PFC, medial PFC, precuneus, and PCC (Fig. 3 - ***Panel C* [bottom]**). It is important to clarify here that, while similar brain regions were seen appearing as key nodes within the different functional networks, the FC patterns were markedly different between the two TMS targets and across different subjects.

### The combined influence of individual cortical geometry and individual connectivity structure on TMS target connectivity patterns

In order to evaluate the similarity of the network engagement in the dlPFC and OFC (when using subject-specific E-fields) between the group average FC matrix and subject-specific FC matrices, we studied the Pearson correlation between the percentage of E-field vertices for the VAN, FPN, and DMN, across all subjects, for these two FC matrix variants. For dlPFC targets, the group average and subject-specific FC matrices showed a higher correlation in the percentage E-field vertices that maximally correlated with the DMN (r=0.54), than the VAN and FPN (r=0.19 and r=0.44, respectively). We found the opposite to be true in the OFC. Here, we observed that there was a higher correlation between the percentage of E-field vertices preferentially correlated with FPN (r=0.40) than with DMN (r=0.23), when comparing group average and subject-specific FC matrices.

Similarly, to determine the similarity of the network engagement in the dlPFC and OFC (when using subject-specific FC matrices) between the fixed (‘Ernie’) E-field and subject-specific E-field, we looked at the Pearson correlation between the percentage of E-field vertices, for the VAN, FPN, and DMN, across all subjects, for the two E-field variants. In the dlPFC, the fixed and subject-specific E-fields showed a higher degree of correlation in the proportion of vertices in each subject’s specific FC matrix maximally targeting the VAN (r=0.80), followed by the FPN (r=0.63), and lastly the (r = 0.56). In the OFC, we noticed the reverse to be true. The fixed and subject-specific E-field showed a higher correlation in the proportion of vertices correlated with the DMN (r=0.4), than with the FPN (r=0.2). A comparison of the differences in patterns of FC between the fixed E-field and the subject-specific E-field can be found in Fig. 4 - ***Panel A***. Taken together, these analyses indicate that the average E-field and group-average FC data are able to predict which networks are targeted, for some networks, but cannot necessarily tell the degree to which each individual network is specifically targeted across subjects.

**Figure 4:**
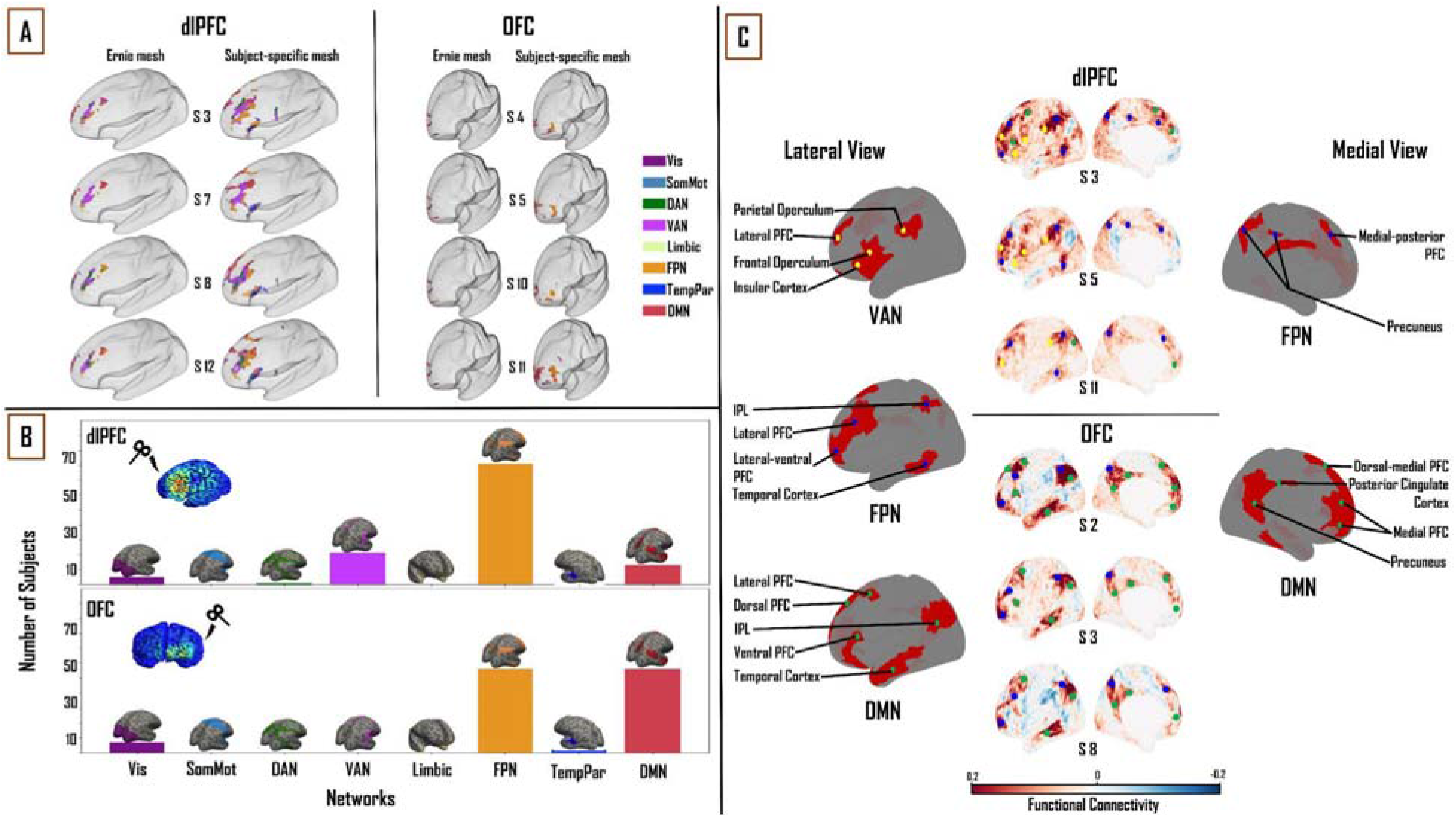
Combined influence of individual cortical geometry and individual FC on TMS target connectivity. **<u>A)</u>** Differences in patterns of FC between fixed (‘ernie’) and subject-specific E-fields. **<u>B)</u>** Like Figures 2, 3 above, summary statistics of network engagement over all 121 subjects shows that the VAN, FPN, and DMN were the main networks engaged from the dlPFC [top]; while the FPN and DMN were the main networks engaged from the OFC [bottom]. **<u>C)</u>** *Top:* lateral and medial views of dlPFC FC maps in subjects 3, 5, and 11, highlighting the key regions that are functionally connected across the VAN, FPN, and DMN. These regions lie mainly in the frontal, parietal and temporal cortices. *Bottom:* lateral and medial views of OFC FC maps in subjects 2, 3, and 8, highlighting the key regions that are functionally connected across FPN and DMN. These regions lie mainly in medial-frontal, cingulate, and posterior parietal cortices. VAN = ventral attention network, FPN = frontoparietal network, DMN = default-mode network.

#### Functional network connectivity based on subject-specific E-fields and subject-specific functional connectivity matrices

Again, one-way ANOVA showed a significant main effect of “NETWORK” for dlPFC (F_(1,7)_ = 267.71, p < 0.0001, η2 = 0.69) and OFC (F_(1,7)_ = 326.36, p < 0.0001, η2 = 0.73). In the dlPFC, VAN (22.2 ± 9.7%), FPN (34.6 ± 10.2%) and DMN (18.4 ± 8.1%) were seen to account for 75% of the E-field vertices. FPN (38.1 ± 12.9%) and DMN (37.8 ± 16.5%) accounted for 76% of the E-field vertices in the OFC. Unlike the previous two sections, the FPN was the most engaged network from the dlPFC, followed by the VAN and DMN across the group (81, 21 and 13 out of 121 subjects, respectively). The DMN and FPN were equally engaged from the OFC (56 subjects each out of 121) (Fig. 4 - ***Panel B***). Similar to the previous sections, pairwise t-tests showed strong interactions between the TMS target areas and the three major networks. We found the OFC target region showed greater connectivity to the DMN (T = 14.2; p <0.0001) and FPN (T = 2.6; p<0.01) relative to the dlPFC target region, whereas the dlPFC target region showed greater connectivity to the VAN (T = 23.3; p <0.0001).

#### The relation between TMS targets and downstream brain regions

At the dlPFC, on the lateral left cortical surface, the IPL, frontal and parietal opercula, and lateral PFC were the major VAN nodes. Lateral FPN nodes included the IPL, the lateral-ventral and lateral PFC, and the posterior region of the middle and inferior temporal gyri. Medially, the FPN nodes consisted of the medial-posterior PFC and the precuneus. The lateral DMN nodes included the lateral PFC, ventral PFC, and IPL. Medially, the DMN nodes included the dorsal-medial and medial PFC. Once again, similar to the previous sections, no VAN nodes were observed on the medial cortical surface (Fig. 4 - ***Panel C* [top]**). At the OFC, on the lateral cortical surface, the FPN nodes included the IPL, lateral-ventral, and lateral PFC. Medial FPN nodes included the precuneus and medial-posterior PFC. Within the DMN, the IPL, lateral and dorsal PFCs, and the anterior region of the middle and inferior temporal gyri were noted as the main lateral nodes. Medially, the precuneus, PCC, dorsal-medial, and medial PFC made up the DMN nodes (Fig. 4 - ***Panel C* [bottom]**).

## Discussion

In this study, we sought to characterize comprehensively two major therapeutic TMS target sites, the dlPFC and the OFC, in terms of a) their patterns of FC to other regions and canonical brain networks, and b) the level and sources of inter-subject variability in those connectivity patterns, using a combination of E-field modelling and analyses of resting-state fMRI data in a group of healthy subjects. With respect to the first of these, our chief conclusion was that three major functional networks were targeted across the dlPFC and OFC: VAN, FPN, and DMN in the dlPFC, and FPN and DMN in the OFC. Furthermore, while these major networks consistently appeared across all subjects, the relative connectivity strengths between the networks, as well as the downstream nodes within each network, varied considerably on a subject-wise basis. This is consistent with previous observations in both animals (Bergmann et. al., 2020) and humans (Mueller et al., 2013). With respect to the question of the level and sources of variability, our approach was to separate, and study both independently and in combination, the effects of variability in skull anatomy and cortical geometry (as encapsulated in subject-specific E-field maps), and of variability in subject-specific FC maps. These analyses showed that the average E-field and group-average FC data are able to predict which networks are targeted, for some networks, but cannot necessarily tell the degree to which each individual network is specifically targeted across subjects. In the following, we discuss the key components of these findings, their interpretation in relation to previous work, and highlight important caveats and limitations.

### Connectivity of TMS targets

Regions showing strong FC with TMS targets give us some insight into the potential functional effects of TMS stimulation. The results of our study revealed the VAN, FPN, and DMN as the major functional networks targeted by dlPFC TMS, and the FPN and DMN as the major networks targeted by OFC TMS. Specific network nodes within each of these networks were observed. Some network nodes such as the lateral PFC (VAN, FPN, DMN) and precuneus (FPN, DMN) were seen across all subjects and in multiple networks. On the other hand, certain networks and nodes were specific to dlPFC TMS or OFC TMS. For example, the connectivity to the VAN is seen in dlPFC TMS, but not in OFC TMS. At the level of individual network nodes, the PCC is a DMN-specific node in E-field FC patterns for OFC TMS targets, but not dlPFC TMS targets. Furthermore, while many similar nodes occur across these networks in multiple subjects, the overall pattern of FC observed varies from subject to subject. The relevance and important functions of the three major functional networks highlighted in these results (FPN, DMN, and VAN) are outlined below.

The FPN is a system implicated in cognitive control for regulating goal-driven behavior. This network is believed to play a key role in problem-solving, as well as actively preserving and editing the information stored in working memory (Uddin et al., 2019). The DMN is active during resting wakefulness when an individual is not actively engaged with external stimuli (Fox et al., 2005). The DMN is also involved in ruminative processes, specifically with thoughts concerning oneself, their past or future events (Andrews-Hanna, 2012). The VAN, sometimes called the salience network, keeps track of salient events (triggered by sensory stimuli) and plays a role in response inhibition or selection (Menon & Uddin, 2010). The VAN is crucial for spontaneous cognitive control, where it helps switch between the DMN’s ruminative/self-reflective functions to the FPN’s task-based/externally driven functions (Menon, 2011; Menon & Uddin, 2010). Neuroimaging studies have shown that the heterogeneous nature of MDD and its subtypes may emerge as a result of unique patterns of disruption in these networks’ dynamics (Feffer et al., 2018). Indeed, multiple research groups have begun to utilize abnormal FC patterns to characterize MDD subtypes (Peng et al., 2012), showing how differences in spontaneous dynamics might potentially lead to different clinical outcomes (Fox et al., 2012). In line with this evidence, our results suggest that a connectivity-based targeting strategy for optimizing network engagement on a per-subject basis may be beneficial for optimizing clinical responses.

### Implications for rTMS therapy

A systematic review of 25 neuroimaging studies of MDD summarized that hypoconnectivity occurs within the FPN and VAN, while regions that were a part of the DMN exhibited hyperconnectivity (Kaiser et al., 2015). There are multiple inhibitory and excitatory rTMS protocols used for inducing region-specific changes in neural activity. Excitatory paradigms include intermittent theta-burst stimulation (iTBS) and high-frequency (10-20Hz) rTMS, whereas prevalent inhibitory paradigms are continuous theta-burst stimulation (cTBS) and low frequency (∼1Hz) rTMS (Downar & Daskalakis, 2013; Huang et al., 2005). The implications from our results may further enhance rTMS targeting practices by informing not only regions but also the type of paradigm to use as well. In our study, a majority of subjects had a higher network engagement with the FPN or VAN than DMN, in the dlPFC (Figure 4 -***Panel B* [top]**). Therefore, one way of understanding the positive therapeutic effects of applying iTBS or high-frequency rTMS at the dlPFC in MDD patients may be that this intervention could result in an excitation - and perhaps renormalization - of the VAN and FPN networks, which show hypoconnectivity in MDD (Kaiser et al., 2015). At the OFC, our results show a similar network engagement between the DMN and the FPN (Figure 4 - ***Panel B* [bottom]**). However, the DMN is engaged more from the OFC than from the dlPFC. Thus, in this case, applying cTBS or low-frequency rTMS may be expected to inhibit the DMN, and again potentially achieve a renormalization of DMN hyperconnectivity in MDD. However, we would like to note here that due to similar network engagement of the FPN and DMN from the OFC, the extent to which one is engaged over the other depends on that subject’s specific FC. A more DMN-centric target could be the dmPFC (Downar & Daskalakis, 2013) as the DMN has more nodes on the medial cortical surface. Targeting specific networks with unique rTMS paradigms in this way may alleviate depressive symptoms more efficiently. In the future, this line of research may be further explored to identify which networks are affected in a given patient, and selectively target them, thereby potentially personalizing rTMS therapy for individuals with MDD.

### Functional Connectivity Variability

We calculated the Pearson correlation coefficient between the percentage of E-field vertices for the VAN, FPN, and DMN, across all subjects, for the group average FC matrix and the subject-specific FC matrix. The group average FC matrices were able to predict inter-individual differences in how networks were targeted from the dlPFC and OFC to a certain extent. In the dlPFC, we observed that the 1003-subject HCP average FC matrix was able to predict what DMN connectivity would be with the subject-specific FC matrices (r=0.54) better than with the VAN (r=0.19) and FPN connectivity (r=0.44). However, this observation was reversed in the OFC. Here, the average FC matrix was able to predict what FPN connectivity would be with subject-specific FC matrices (r=0.40) better than with the DMN (r=0.23). One explanation for this finding is that DMN has a more consistent spatial pattern across subjects than the VAN or FPN, such that subject-level and group-level patterns are relatively more similar than for other networks. However, this line of reasoning does not explain why a pattern reversal occurs at the OFC. In summary, our results confirm the general intuition that using an average FC matrix provides a gross estimate of what the targeted networks might be, but precise targeting requires each subject’s specific FC data.

### E-field Variability

We observed the mean subject-specific thresholded E-field size to be 54.3 and 16.2 mm^2^, in the dlPFC and OFC, respectively. However, the E-field size varied considerably across subjects, with a standard deviation of ± 18 mm^2^ in the dlPFC and ± 8 mm^2^ in the OFC. This high intersubject variability of dlPFC and OFC E-fields can be attributed to variability in subject-specific cortical geometry. Consistent with this, the boundaries between the five main tissue types have been shown to affect E-field distributions. These include the skin, skull, CSF, white matter, and gray matter (Thielscher et al., 2011), and are highly variable across subjects. Furthermore, this E-field variability had a knock-on effect on variability in the connectivity of the dlPFC and OFC stimulation targets to downstream functional networks. In the dlPFC, the normative template E-field (from the ‘ernie’ brain) and subject-specific E-fields showed the highest network engagement correlation with the VAN (r=0.80), followed by the and FPN (r=0.63) and then the DMN (r=0.56), in each subject’s specific FC matrix. In the OFC, the opposite was found to be true. The normative template E-field and subject-specific E-field showed a higher network engagement correlation with the DMN (r=0.4), than the FPN (r=0.2). A potential reason for this observation is the large difference in size between the template E-field and the subject-specific E-fields. In the dlPFC, the subject-specific E-fields are nearly twice as large (mean = 54.3 mm^2^) as the template E-field (31 mm^2^). The difference is greater in the OFC, with the subject-specific E-fields (mean = 16.2 mm^2^) being over two and half times the size of the template E-field (6.3 mm^2^). The additional vertices in each subjects’ specific E-field tend to target the VAN and FPN in the dlPFC, as the E-field vertices here are predominantly present on the ventral/lateral surface of the prefrontal cortex. The DMN, being more medially located overall, therefore has lower connectivity to dlPFC when the template E-field is used than when subject-specific E-fields are used (Figure 4 - ***Panel A* [left]**). Furthermore, this line of reasoning can be extended to account for the pattern reversal observed in the OFC, where the template E-field does not account for the additional vertices in the subject-specific E-fields which are spread more laterally, targeting the FPN (Figure 4 -***Panel A* [right]**), and hence shows a pattern targeting the DMN but not the FPN.

### Caveats and Limitations

While the results of this study are promising, there are some important caveats and limitations to highlight.

One important limitation is the fact that we only use the left hemisphere to study TMS target connectivity. The reason for this choice was in part practical (simplifying surface-based analysis), but also reflected the fact that as a rule, we expect FC patterns to the two TMS target zones to be dominated by intrahemispheric connections, with the obvious exception of the contralateral homologue (i.e., right dlPFC and right OFC). By using FC data from only one hemisphere, we are therefore potentially missing some important differences between subjects and TMS targets in their connectivity to the contralateral homologues. Moreover, our choice was also driven by common practise in clinical settings which use the left hemisphere as a target for rTMS treatments. However, given that our focus here is on patterns of FC to distal cortical regions that are outside of either the primary target area or its hemispheric homologue, we feel this approach is justified.

Another important limitation is the E-field threshold and its effect on resultant FC calculations. In this study, the E-field threshold was set to 0.9 V/m, which is slightly lower than that used by (Romero et al., 2019). Our justification for this choice is that higher thresholds (i.e., above 0.9 V/m) shrink the E-field sizes, especially in the OFC, and hamper FC calculations. The problem remains however that in the field of TMS more broadly, it is not yet clear what a ‘correct’ E-field threshold should be. Thus, we explored the effects of using a range of E-field values and these results are detailed in the ***Supplementary Information*** section (*below*). Often the ‘correct’ E-field threshold is conceived as the minimum induced current necessary to depolarize neuronal membranes and cause them to fire. Subthreshold effects (i.e. ones not resulting from action potential induction at the primary stimulation site) may nevertheless potentially have an important role in TMS responses; for example by facilitating the occurrence and frequency of suprathreshold events. Spatially, the question of E-field thresholding relates quite closely to the question of E-Field size and extent (since high thresholds usually ‘trim’ the edges of activated areas, eliminating vertices around the penumbra first).

A further, related, limitation is that interpretation of our results, and those from related work (e.g. Opitz et al., 2016) rests heavily on the notion that FC can serve as a reliable indicator of which downstream brain regions, distal to the TMS target site, would themselves be ‘activated’, or otherwise affected, by TMS administration. In defense of this principle, multiple studies have shown experimentally that neuronal activation as a result of TMS is not limited to the cortical circuits closest to the scalp (Bergmann et al., 2021; Hawco et al., 2018; Siebner et al., 2009; Solomon-Harris et al.,, 2016). These studies show that initial local neuronal activation spreads across cortical and subcortical regions to neighboring and distant parts of the brain. In other words, it appears to be impossible to stimulate a single region of the brain with TMS without affecting a large number of downstream network nodes. While further studies are required to decipher the precise pathways taken to activate these downstream nodes, FC maps offer a plausible proxy for assessing which networks are being engaged for a given TMS target region.

Importantly, the subjects chosen for this study are from a normative, healthy sample (from the HCP database). However, MDD patients may have different/altered connectivity patterns that the healthy subject patterns may not be representative of. While previous research has looked at connectivity-based targeting in MDD patients with promising results (Weigand et al., 2018), a full-fledged clinical trial evaluating this method is yet to be undertaken (Cash et al., 2020).

One potential improvement to the methodology used here that may be considered for future work is to evaluate alternative TMS coil options. Here, we have chosen to use the Magstim 70mm Figure-8 coil to run our TMS simulations in SimNIBS. Used in both clinical and research settings, it has been shown that Figure-8 coils allow for a more focused stimulation of the target site (Thielscher & Kammer, 2004) than other design options. We used the Figure-8 coil type to run our simulations for both the dlPFC and OFC. However, the thresholded E-field size difference between these two TMS targets in our analyses is likely due mainly to their anatomical locations, and distance from the stimulating coil. The dlPFC is located at the frontal lobe and lies on the lateral and dorsal surface of the medial convexity, fairly close to the scalp surface. The OFC, on the other hand, is a large gray matter shelf located on the ventral surface of the frontal lobes, above the orbit of the skull. As a result, a large portion of the OFC is not accessible via the Fp1 electrode position on the scalp (which is more frontal in location than ventral). Thus, a typical Figure-8 coil cannot target the OFC as effectively as it can the dlPFC, owing to the inconsistency of the targeting surface. To address this issue, alternate coil designs have been proposed, such as crown-shaped coils, C-shaped coils (Deng, Peterchev, & Lisanby, 2008), and H-shaped coils (Levkovitz et al., 2009), which have been developed to target deeper cortical regions. It will be valuable to analyze the resulting E-fields produced by these coils with our current methodology, to better establish the effects of TMS with all potentially available coil configurations on novel treatment sites, such as the ventral OFC and regions of the medial PFC.

### Conclusions and Future Directions

We have presented data characterizing the FC patterns of canonical therapeutic TMS targets and the key dimensions of their variability across subjects. Our results show that the VAN, FPN, and DMN are the major functional networks targeted by dlPFC TMS, and the FPN and DMN are the major networks targeted by OFC TMS. These results provide important insights into the functional neuroanatomical effects of clinical TMS stimulation protocols. Importantly, our results validate the general intuition that using a normative group-averaged FC matrix provides only a coarse estimate of what the targeted networks might be, and precise targeting requires each subject’s specific FC data.

Our hope is that these insights prove useful as part of the broader effort by the psychiatry, neurology, and neuroimaging communities to help improve and refine TMS therapy, through a better understanding of the technology and its neurophysiological effects. Further work shall be needed to evaluate the predictive and clinical utility of the TMS target fMRI FC profiles, through both prospective and retrospective clinical neuroimaging studies in MDD patients. Progress on the neurobiological question of what are the network-level effects of TMS stimulation, however, necessitates an integrative approach combining various neuroimaging and physiological modalities, and various quantitative techniques. In particular, characterization of the structural connectivity between TMS targets and their downstream networks using diffusion-weighted MRI tractography analyses, which can serve as a useful proxy for axonal connectivity between various brain regions, shall be an important area of investigation that should complement the results reported in the present study. How do target region connectivity profiles from tractography connectivity compare to their FC analogs? How should discrepancies and convergences between structure and function be interpreted in relation to expected TMS effects? Another important question we hope to answer in the future is how using different locations with the same target regions affects downstream FC. For example, both the F3 and F5 electrode locations fall within the dlPFC. It would be interesting to study the differences in network engagement in this situation. Ultimately the best-known general strategy for reconciling such questions (and one that we are currently pursuing intensively) is to develop validated and predictively accurate computational models of brain stimulation responses, that include relevant biological detail but are also sufficiently scalable to allow whole-brain activity simulations. In future work, our aim is to use mechanistic modelling approaches to formalize and test hypotheses around synaptic-, local circuit-, and network-level mechanisms in brains receiving noninvasive stimulation, and to use the insights obtained to help improve the efficacy of TMS in the clinic.

## Acknowledgments

We are grateful to the Krembil Foundation, CAMH Discovery Fund, and Labatt Family Network for the generous funding support that has made this research possible. *CRediT author contributions:* SH: Conceptualization, Methodology, Formal analysis, Writing - Original Draft, Writing - Review & Editing, Visualization; JDG: Conceptualization, Methodology, Writing - Review & Editing, Supervision, Funding acquisition; DM: Writing - Review & Editing, Visualization; FM: Writing - Review & Editing.

## Supplemental Information

### Effect of varying E-field thresholds on the network engagement of TMS targets

We use E-field thresholds in order to determine how much of the underlying cortical tissue is being activated by TMS stimulation. This is something that is not clearly understood, and moreover, the E-field threshold can vary across individuals (due to unique cortical geometry) and can also vary based on the experimental paradigm that is being used (excitatory vs. inhibitory TMS). Our default E-field threshold choice was 0.9 V/m, for reasons explained in the *Discussion*. However, in order to better understand the stability of our results across a range of potential threshold values, we further investigated the network engagement of TMS targets over a specific set of E-field thresholds ranging from 0.7 V/m to 0.9 V/m in increments of 0.05 V/m for a total of 5 different potential E-field thresholds. Our results showcase the effects of larger E-fields on the network engagement of TMS targets as expected, and are highlighted below.

### E-field size and distribution

At the original threshold of 0.9 V/m, the E-field size ranged from 120 to 1125 vertices (mean =542.9 ± 179.8) in the dlPFC and from 32 to 607 vertices (mean = 161.8 ± 80.5) in the OFC. As noted in the *Methods*, E-field sizes are reported in terms of ‘number of vertices’, where the average area of the face associated with vertex triplets is 0.05 mm^2^. As we lowered the E-field threshold, the E-field sizes were seen to increase in both the dlPFC and OFC. At the lowest threshold value we looked at (0.7 V/m), the dlPFC E-field size ranged from 455 to 2370 vertices (mean = 1241.7 ± 329). The OFC E-field size ranged from 186 to 1301 vertices (mean = 444.7 ± 147.9). Furthermore, in both the dlPFC and OFC, lowering the E-field threshold led to the spatial distribution of the respective E-fields reaching other areas of the cortex that were not the intended TMS target, and are unlikely to receive supra-threshold stimulation. For example, in the dlPFC, step-wise lowering of the E-field threshold from 0.9 V/m to 0.7 V/m saw the inclusion of parts of the motor and somatosensory cortex. On the other hand, in the OFC, lowering the E-field threshold led to the inclusion of parts of the dlPFC as part of the OFC E-field. These differences are showcased in ***Supplemental Figure 1***.

**Supplemental Figure 1:**
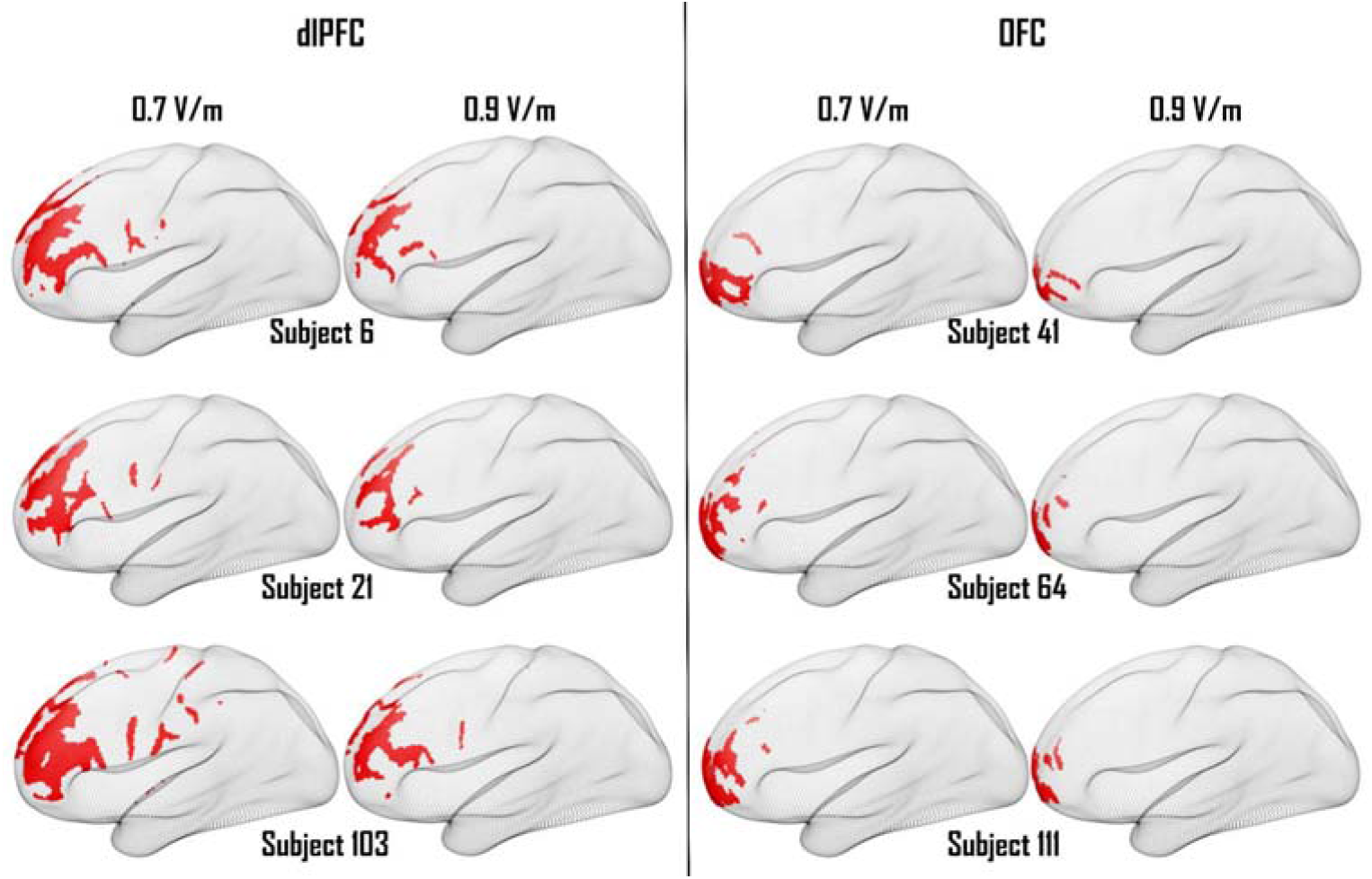
The difference in the size and spatial extent of dlPFC and OFC E-fields are seen at the lowest (0.7 V/m) and highest (0.9 V/m) E-field thresholds.

### Network engagement of TMS targets across a range of E-field thresholds

We observed that our results were highly stable across the various E-field thresholds, with one interesting exception.

The VAN, FPN, and DMN were seen as the major networks engaged from the dlPFC. The VAN engagement was seen to increase as the E-field threshold was increased. At the lowest E-field threshold (0.7 V/m), the VAN showed the highest engagement of all the networks in only 5 subjects (∼4%). However, this number was seen to increase to 21 subjects (∼17%) as the E-field threshold was increased to 0.9 V/m. We observed that the FPN was the most engaged network from the dlPFC in the majority of subjects, regardless of the E-field threshold value. At a threshold of 0.7 V/m, 92 of the 121 (∼76%) subjects had the highest network engagement with the FPN. As the threshold was increased to 0.9 V/m, this number dropped to 81 (∼67%). The DMN network engagement was relatively consistent across the different E-field thresholds. Maximum DMN engagement ranged from 16 subjects (∼13%) at 0.7 V/m to 13 subjects (∼10%) at 0.9 V/m (***Supplemental Figure 2A***).

In the OFC, we noticed an interesting pattern of network engagement between the FPN and DMN. As we increased the E-field threshold from 0.7 V/m to 0.9 V/m, there was a steady decrease in FPN engagement and a steady increase in DMN engagement. At 0.7 V/m, the FPN was the maximally engaged network in 91 of 121 subjects (∼75%), but at 0.9 V/m, this value dropped to 56 subjects (∼46%). On the other hand, maximal DMN engagement was seen in 24 subjects (∼20%) at 0.7 V/m but this number increased to 56 subjects (∼46%) at 0.9 V/m matching the value for the FPN. Thus, the maximally engaged network (FPN vs. DMN) for OFC stimulation depends on the E-field threshold level (***Supplemental Figure 2B***).

**Supplemental Figure 2:**
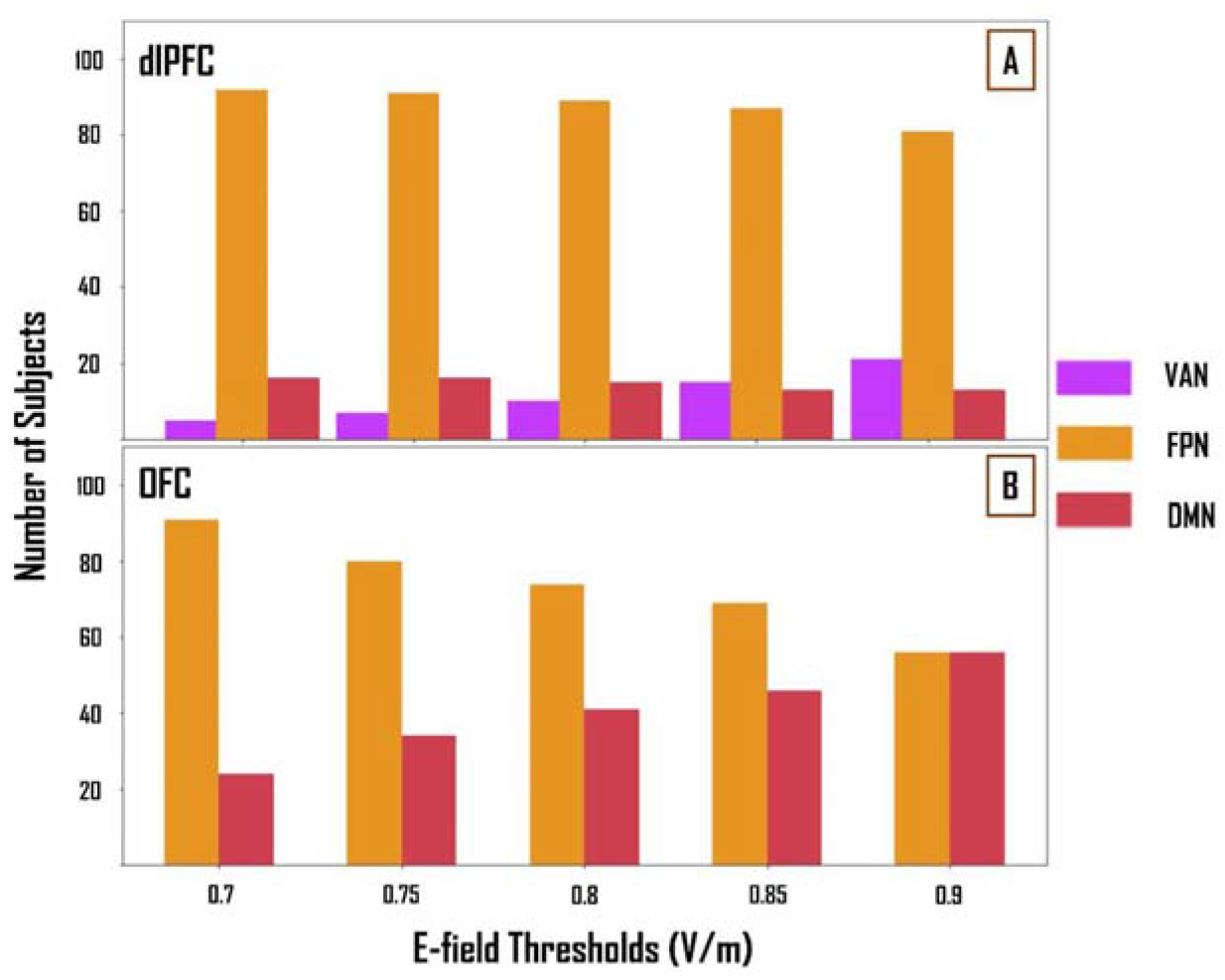
The network engagement of the VAN, FPN, and DMN are shown across multiple E-field thresholds across subjects. In the dlPFC, the FPN is the most engaged network across a majority of subjects regardless of the E-field threshold. In the OFC, the FPN engagement decreases, and the DMN engagement increase as the E-field threshold is increased, across subjects.

### Implication of findings

In addition to the main analysis of this paper, we looked at the effect of varying E-field thresholds on the network engagement of TMS targets. As expected, the overall E-field size and distribution were inversely proportional to the E-field threshold.

In the dlPFC, we observed that at lower thresholds, the E-fields included regions that were not the intended TMS target like the primary motor and somatosensory cortices. However, this didn’t affect the overall network engagement drastically. The FPN was the most engaged functional network in a majority of subjects regardless of the E-field threshold. By increasing the E-field threshold (i.e., reducing the E-field size), we noticed that the VAN saw a slight increase as the most engaged network in some subjects. This could be due to the elimination of vertices on the more lateral/ventral surface of the PFC which are mainly associated with the FPN. The DMN was not the maximally engaged network across most subjects as the dlPFC E-field primary lies on the lateral/dorsal surface of the PFC while the DMN nodes are mainly more medial in location.

In the OFC, we observed an interesting relationship between the E-field threshold and corresponding network engagement of the FPN and DMN. As the E-field threshold was increased, we report that the FPN engagement was reduced while the DMN engagement was increased. The reason for this observation is evidenced directly from the E-fields themselves. At lower E-field thresholds (larger E-fields), the vertices of the E-field include regions of the dlPFC as part of the OFC E-field. As seen in our results above, dlPFC E-fields tend to maximally engage the FPN in a majority of the subjects. Therefore, this spill-over of E-field vertices from the OFC E-field leads to the inclusion of the dlPFC areas and thus a higher FPN engagement. Conversely, by increasing the E-field threshold (smaller E-fields), these additional dlPFC vertices are trimmed and this leads to a lower FPN engagement and a higher DMN engagement.

As noted in the main manuscript, we would again like to highlight the fact that our results show equal targeting of the FPN and DMN from the OFC at our original threshold (0.9 V/m). However, the DMN is engaged in a higher number of subjects from the OFC, than from the dlPFC, but the extent to which each network is engaged over the other comes down to the individual’s FC.

